# Regulating expression of mistranslating tRNAs by readthrough RNA polymerase II transcription

**DOI:** 10.1101/2021.09.16.460584

**Authors:** Matthew D. Berg, Joshua R. Isaacson, Ecaterina Cozma, Julie Genereaux, Patrick Lajoie, Judit Villén, Christopher J. Brandl

## Abstract

Transfer RNA (tRNA) variants that alter the genetic code increase protein diversity and have many applications in synthetic biology. Since the tRNA variants can cause a loss of proteostasis, regulating their expression is necessary to achieve high levels of novel protein. Mechanisms to positively regulate transcription with exogenous activator proteins like those often used to regulate RNA polymerase II (RNAP II) transcribed genes are not applicable to tRNAs as their expression by RNA polymerase III requires elements internal to the tRNA. Here, we show that tRNA expression is repressed by overlapping transcription from an adjacent RNAP II promoter. Regulating the expression of the RNAP II promoter allows inverse regulation of the tRNA. Placing either Gal4 or TetR-VP16 activated promoters downstream of a mistranslating tRNA^Ser^ variant that mis-incorporates serine at proline codons in *Saccharomyces cerevisiae* allows mistranslation at a level not otherwise possible because of the toxicity of the unregulated tRNA. Using this inducible tRNA system, we explore the proteotoxic effects of mistranslation on yeast cells. High levels of mistranslation cause cells to arrest in G1 phase. These cells are impermeable to propidium iodide, yet growth is not restored upon repressing tRNA expression. High levels of mistranslation increase cell size and alter cell morphology. This regulatable tRNA expression system can be applied to study how native tRNAs and tRNA variants affect the proteome and other biological processes. Variations of this inducible tRNA system should be applicable to other eukaryotic cell types.

## INTRODUCTION

Transfer RNAs (tRNAs) are the vehicle that bring amino acids to the growing polypeptide chain at the ribosome and read the three base codons that define protein sequence^1–4^. Both as intact molecules and as fragments, tRNAs play other essential roles in cell physiology (reviewed in [5]). Manipulating tRNA function has numerous applications in synthetic biology and genetic code expansion, as well as in understanding genetic code evolution and the mechanisms cells use to maintain proteostasis.

One application for tRNA variants is in generating mistranslation. Mistranslation occurs when an amino acid that differs from what is specified by the “standard” genetic code is inserted into a protein during synthesis. Mistranslation occurs naturally in all cells at frequencies ranging from 1 in 3000 to 1 in 10^6^ depending on the amino acid (reviewed in [6]) with higher levels occurring in response to various environmental conditions (see for example [7–10]). Excessive mistranslation leads to proteotoxic stress and slows cell growth^11–13^; however, cells use protein quality control mechanisms to tolerate mistranslation at levels approaching 8-10%^14–16^. Proteins arising from ambiguous translation caused by mistranslating tRNAs are called statistical proteins and have the potential for a broader range of function than a homogeneous protein^17^. Statistical proteins have regulatory functions in cells^18^ and applications in synthetic biology and biotechnology by increasing functional diversity^19^.

A number of systems have been developed to regulate expression of RNA polymerase II (RNAP II) transcribed mRNA encoding genes. These often involve placing transcriptional activator binding sites into 5’ promoter sequences, which then allow the gene to be regulated by a heterologous transcriptional activator protein. A classic example of this uses the Gal4 activator protein and its binding sites to regulate a specific gene. In contrast to mRNAs, in eukaryotic cells tRNAs are transcribed by RNA polymerase III^20^ (RNAP III). The key elements for RNAP III recruitment are the A box and B box^21–23^. These elements are internal to the tRNA encoding region and thus cannot be modified without altering the tRNA structure. We have tried to block RNAP III transcription by inserting tet operator sites within tRNA intron sequences with minimal success. We do note that the Marschalek and Capone labs had some success repressing tRNA expression by placing tet or lac operator sites upstream of a tRNA gene^24–26^, while Herschbach and Johnson (1993)^27^ found the yeast α2 operator/repressor could not repress RNAP III genes. Other strategies that do not rely on transcriptional control have been used to regulate tRNA levels. One approach, developed by Zimmerman *et al*. (2018)^16^, is to regulate tRNA turnover through the rapid tRNA decay (RTD) pathway. The pathway is controlled through a conditional allele of *MET22*, whose substrate inhibits the endonucleases involved in tRNA decay. Disadvantages of regulation through the RTD pathway include that endogenous tRNAs may be affected, tRNA sequence needs to be altered to make them susceptible to decay and not all tRNAs are targeted by this pathway. Similarly, we have expressed otherwise toxic mistranslating tRNAs by altering bases to destabilize tRNA structure, which in turn makes them more prone to decay^15,28^.

Our goal was to develop a routinely applicable system to regulate expression of “non-native” tRNAs in yeast cells for use in synthetic biology applications and to study the impact of high levels of mistranslation. Our approach was based on the observation of Martens *et al*. that intergenic transcription impedes the expression of an adjacent RNAP II transcribed gene^29^. Other work has suggested that transcription through RNAP III regulatory sequences would also inhibit their transcription^30^. In this work we show that tRNA expression is efficiently repressed by placing an RNAP II promoter downstream of a tRNA encoding gene. We use this system to regulate mistranslation in yeast cells, showing that inducing mistranslation causes growth arrest and leads to an accumulation of cells in G1. This system has applications for regulating native and synthetic tRNAs in other eukaryotic systems.

## RESULTS AND DISCUSSION

### RNA polymerase II readthrough transcription regulates tRNA expression

Our goal was to engineer an inducible system to regulated tRNA expression to determine the impact of high levels of mistranslation on cells, including levels that are otherwise toxic. We hypothesized that transcription from an RNAP II promoter through the tRNA encoding gene would interfere with RNAP III transcription and repress tRNA expression. By making the RNAP II promoter inducible, the tRNA would come under inverse regulation.

Serylation of tRNA^Ser^ requires sequences within its long variable arm (Figure 1A) and not within the anticodon^31^. Changing the tRNA^Ser^ anticodon results in serine mistranslation at the codons recognized by the new anticodon. We have demonstrated this for tRNA^Ser^_UGA_, where altering the anticodon to UGG results in the incorporation of serine at proline codons^15,28^. In the context of an otherwise wild type tRNA^Ser^, constructs containing the UGG anticodon can not be transformed into yeast, presumably because their expression causes loss of viability. However, constructs expressing tRNA^Ser^ with a UGG anticodon can be transformed into yeast if they contain a secondary mutation, for example G26A, that reduces their steady state level^28^. Strains expressing tRNA^Ser^_UGG,G26A_ introduced on a centromeric plasmid mistranslate serine at proline codons at a frequency of approximately 5% and cell growth rate decreases by ∼30% as compared to that seen with wild type tRNA^Ser^_UGA_^15^. We use the effects of tRNA^Ser^_UGG,G26A_ on growth as an initial proxy for mistranslation, as decreased growth correlates with increased mistranslation^15^.

**Figure 1.**
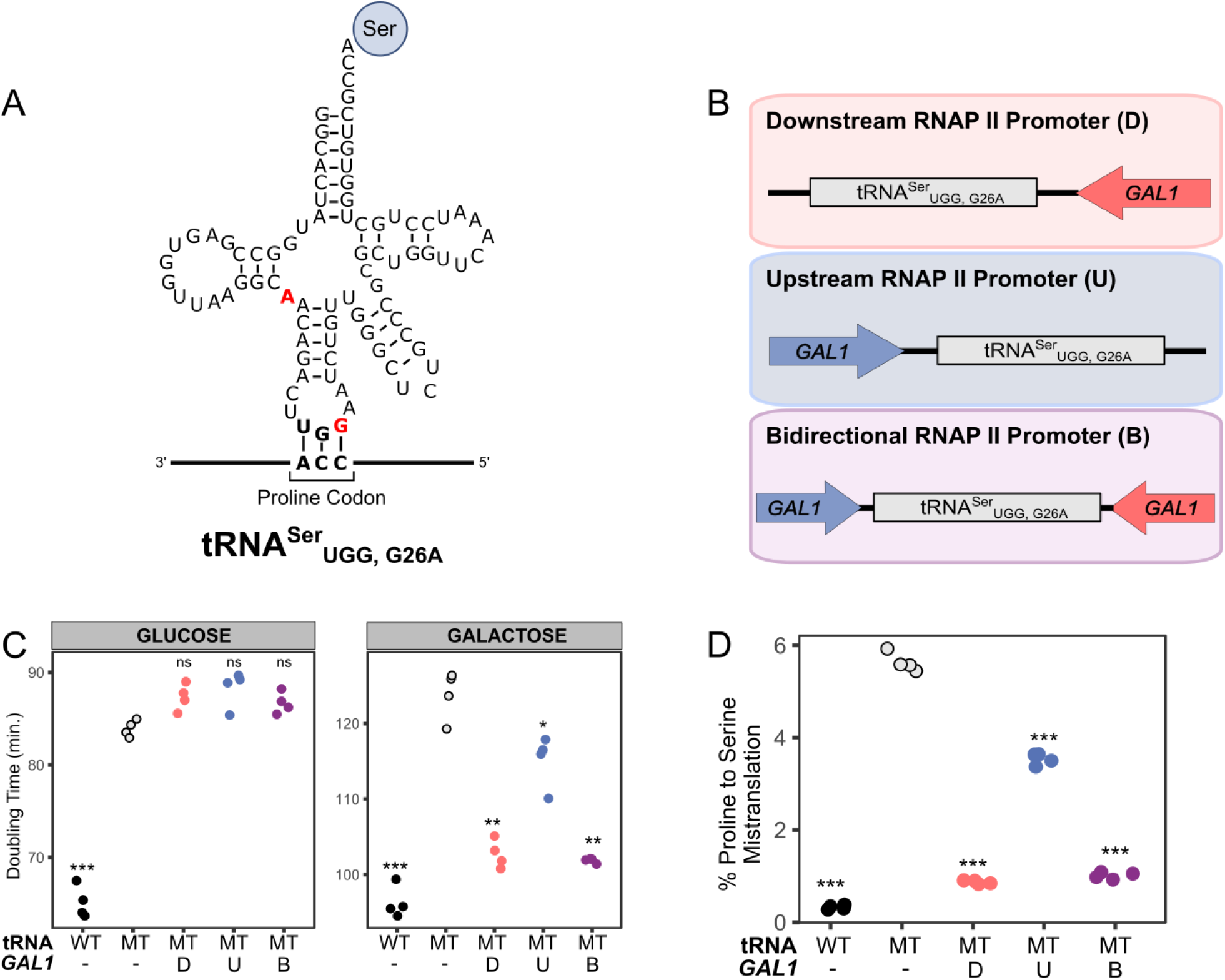
Transcription from a flanking RNAP II promoter decreases mistranslation derived from a mistranslating tRNA gene. **(A)** The structure of tRNA^Ser^_UGG,G26A_. Bases colored red were mutated to allow for non-lethal levels of serine mis-incorporation at proline codons. **(B)** The *GAL1* promoter was cloned either 300 bp downstream (D), 300 bp upstream (U) or both up and downstream (B) of the gene encoding tRNA^Ser^_UGG,G26A_. The sequence directly flanking the tRNA encoding sequence is from *SUP17*. **(C)** Wild type yeast strain BY4742 containing either wild type tRNA^Ser^ (WT), mistranslating tRNA^Ser^_UGG,G26A_ (MT) or the mistranslating tRNA^Ser^_UGG,G26A_ with *GAL1* promoters flanking were grown to saturation overnight in medium lacking uracil. Strains were diluted to an OD_600_ of 0.1 in the same media containing either galactose or glucose as the carbon source and grown for 24 hours at 30°C with agitation. OD_600_ was measured every 15 minutes and doubling time in minutes calculated from the growth curves. **(D)** Mass spectrometry analysis of the cellular proteome was performed on the strains described in (C) grown in galactose to determine the frequency of proline to serine substitution. In each panel, points represent biological replicates and stars indicate significant differences compared to the unregulated tRNA^Ser^_UGG,G26A_ mistranslating strain (Welch’s T-test; * *p* < 0.01, ** *p* < 0.001, *** *p* < 0.0001; ns – not statistically different).

We engineered three constructs to test the regulation of tRNA with a flanking galactose inducible promoter (Figure 1B): one positions the *GAL1* promoter, containing four binding sites for the Gal4 transcriptional activator, 300 base pairs downstream of tRNA^Ser^_UGG,G26A_, another with the *GAL1* promoter 300 bases upstream of tRNA^Ser^_UGG,G26A_ and a third construct with two copies of the *GAL1* promoter, one placed upstream and the other placed downstream of the tRNA. The *GAL1* promoter is highly expressed in medium containing galactose and repressed in medium containing glucose (reviewed in [32]). The centromeric plasmids containing the tRNA and *GAL1* promoter were transformed into a wild type yeast strain and doubling time determined in liquid growth assays. When the strains containing the mistranslating tRNA with flanking *GAL1* promoter constructs were grown in glucose containing medium, where the *GAL1* promoter is inactive, the increase in doubling time was similar to unregulated tRNA^Ser^_UGG,G26A_ (Figure 1C). In contrast, when grown in galactose, where the *GAL1* promoter is active, strains containing the tRNA^Ser^_UGG,G26A_ construct flanked with downstream *GAL1* promoter or flanked with *GAL1* on both sides had doubling times that approached the strain expressing wild type tRNA^Ser^. The strain expressing tRNA^Ser^_UGG,G26A_ with upstream *GAL1* promoter had a doubling time that was intermediate between the strains expressing wild type tRNA^Ser^ and unregulated tRNA^Ser^_UGG,G26A_.

To confirm that mistranslation was inhibited when strains were grown in galactose, we determined mistranslation frequencies in strains containing the regulated tRNA using mass spectrometry. Mistranslation frequency was calculated from the ratio of mistranslated serine peptides identified compared to peptides containing the wild type proline residue. The observed mistranslation frequencies in each strain mirrored the results seen from strain doubling times (Figure 1D). In the strain expressing a wild type serine tRNA, we detected 0.3% substitution of serine at proline codons, whereas in the unregulated mistranslating strain expression tRNA^Ser^_UGG,G26A_ mistranslation frequency was 5.6%. In the strains expressing the mistranslating tRNA with the downstream *GAL1* promoter or with *GAL1* promoter flanking both sides of the tRNA, mistranslation was reduced to ∼ 1%. The *GAL1* promoter positioned upstream also repressed mistranslation, but to a lesser extent (3.5%). These results suggest that transcription from the *GAL1* promoter inhibits the expression of the otherwise toxic tRNA, with the downstream promoter being more effective than the upstream.

To determine if the reduced toxicity of the mistranslating tRNAs was due to Gal4 activity and not another element in the promoter, we replaced the downstream *GAL1* promoter in tRNA^Ser^_UGG,G26A_ with a synthetic *HIS3* promoter containing five Gal4 binding sites^33^ (*HIS3*^5xGAL4^). Again, when grown in glucose containing medium the doubling time of a strain containing this tRNA was similar to one containing unregulated tRNA^Ser^_UGG,G26A_ (Figure 2A). In galactose containing medium, the strain expressing tRNA^Ser^_UGG,G26A_ with downstream *HIS3*^5xGAL4^ grew similarly to the strain containing wild type tRNA^Ser^, supporting the conclusion that Gal4 activity is responsible for inhibiting mistranslation.

**Figure 2.**
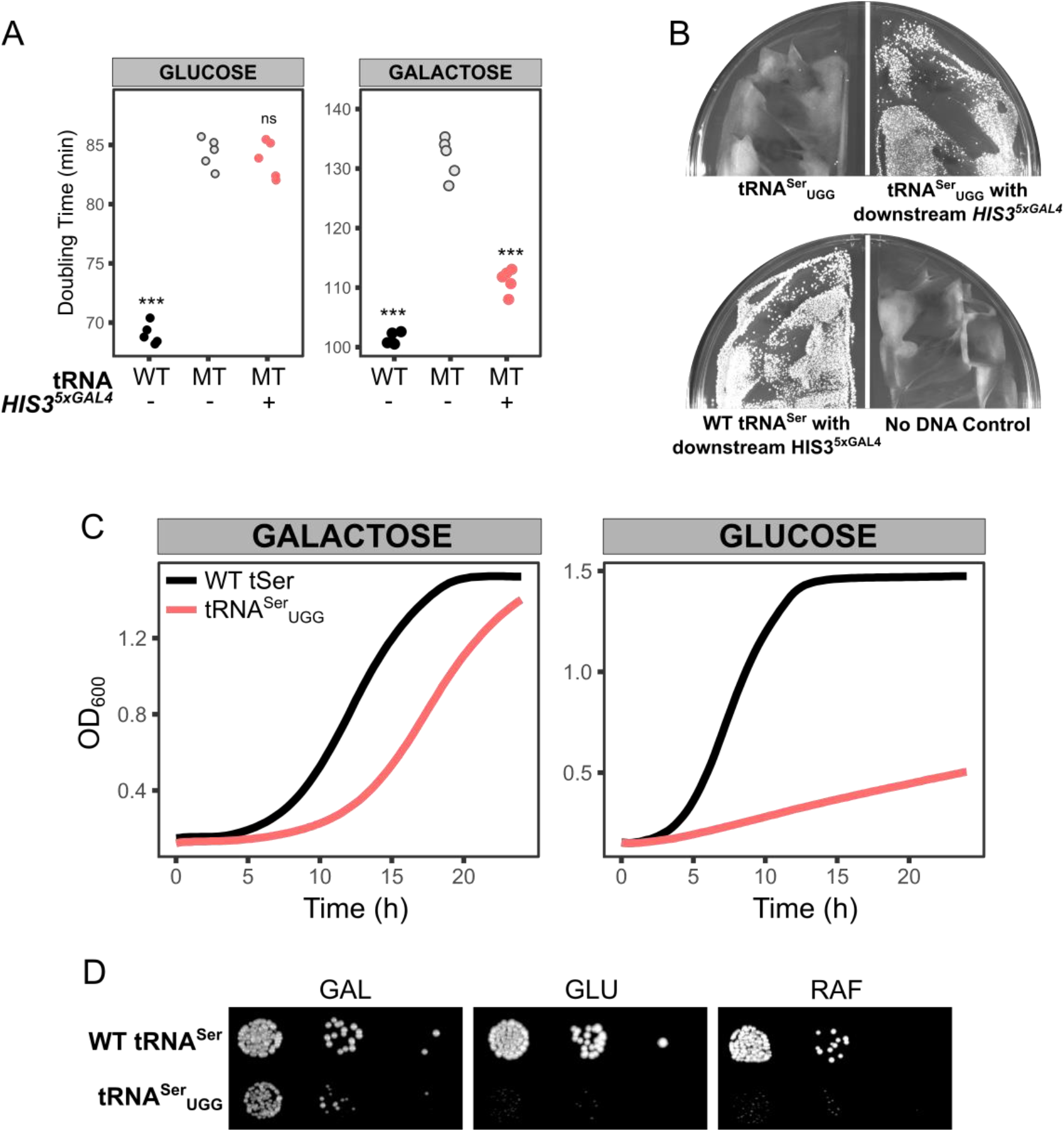
Regulated tRNA expression allows transformation of a tRNA that mistranslates at otherwise lethal levels. **(A)** Wild type yeast strain BY4742 containing centromeric plasmids with either wild type tRNA^Ser^ (WT), mistranslating tRNA^Ser^_UGG,G26A_ (MT) or the mistranslating tRNA^Ser^_UGG, G26A_ with a synthetic galactose inducible promoter containing five Gal4 binding sites (*HIS3*^*5xGAL4*^) were grown to saturation overnight in medium lacking uracil. Strains were diluted to an OD_600_ of 0.1 in the same media containing either glucose or galactose as the carbon source and grown for 24 hours at 30°C with agitation. OD_600_ was measured every 15 minutes and doubling time in minutes calculated from the growth curves. Points represent biological replicates and stars indicate significant differences compared to the unregulated tRNA^Ser^_UGG,G26A_ mistranslating strain (Welch’s T-test; *** *p* < 0.0001; ns – not statistically different). **(B)** Wild type yeast strain BY4742 was transformed with either a *URA3* centromeric plasmid containing wild type tRNA^Ser^ or mistranslating tRNA^Ser^ with UGG anticodon either with or without downstream *HIS3*^*5xGAL4*^ promoter and plated on medium lacking uracil containing galactose. Plates were grown for 3 days at 30°C before imaging. **(C)** Strains from (B) were grown in medium lacking uracil and containing galactose to saturation, diluted to an OD_600_ of 0.1 in medium lacking uracil and containing either glucose or galactose and grown for 24 hours at 30°C with agitation. OD_600_ was measured every 15 minutes. One representative growth curve is shown for each condition. **(D)** Strains from (B) were grown in medium lacking uracil and containing galactose to saturation and plated in 10-fold serial dilutions on media lacking uracil and containing either galactose (GAL), glucose (GLU) or raffinose (RAF). Plates were imaged after two days of growth for the glucose plate and three days of growth for the galactose and raffinose plates.

Next, we determined if this system allows expression of tRNA^Ser^_UGG_ lacking a secondary mutation to dampen function. When unregulated, this tRNA can not be introduced into cells^28^. The *HIS3*^*5xGAL4*^ promoter was cloned downstream of the tRNA^Ser^_UGG_ gene and transformed into yeast. In contrast to unregulated tRNA^Ser^_UGG_, transformants containing the Gal4-regulated tRNA^Ser^_UGG_ were obtained when cells were plated on medium containing galactose (Figure 2B). Growth curves were performed in glucose and galactose media to measure toxicity of tRNA^Ser^_UGG_ in expressed and repressed conditions, respectively (representative growth curves are shown in Figure 2C). In glucose medium, the virtual lack of growth for strains containing tRNA^Ser^_UGG_ demonstrates that the mistranslating tRNA is expressed and toxic to cell growth. Growth of tRNA^Ser^_UGG_ in galactose medium demonstrates that Gal4-mediated transcription limits expression of the otherwise toxic tRNA. Compared to a wild type tRNA^Ser^, there is some toxicity from tRNA^Ser^_UGG_ grown in galactose, likely reflecting low levels of RNAP III mediated transcription of tRNA^Ser^_UGG_ in the repressed state.

To confirm Gal4 driven transcription is required to repress tRNA expression, we assayed growth of the strain containing tRNA^Ser^_UGG_ with the downstream *HIS3*^*5xGAL4*^ in raffinose medium. With raffinose as the carbon source, Gal4 is bound to its binding site but its activity is repressed by Gal80^34^. When plated on medium containing raffinose, the strain expressing tRNA^Ser^_UGG_ with downstream *HIS3*^5xGAL4^ does not grow, indicating that Gal4 driven transcription is essential to repress transcription of tRNA^Ser^_UGG_ (Figure 2D). In agreement with this, overexpressing just the Gal4 DNA binding domain in a *gal4Δ gal80Δ* strain does not suppress tRNA^Ser^_UGG, G26A_ toxicity (Figure S1).

We tested if the distance between the 3’ end of the tRNA and the regulated promoter affected tRNA expression. When cells were grown in galactose medium, the doubling time of strains with the *HIS3*^*5xGAL4*^ promoter 100 bp or 200 bp downstream from the tRNA was not statistically different from when the regulated promoter was placed 300 bp downstream (Figure S2).

To provide a titratable system that did not require switching carbon sources, we placed a minimal *CYC1* promoter with seven TetO binding sites 300 bp downstream of tRNA^Ser^_UGG_ or tRNA^Ser^_UGG,G26A_ (Figure 3A). These constructs and a similar construct containing wild type tRNA^Ser^ were introduced into a strain containing the Tet-Off regulation system^35^. In these strains, the strong TetR-VP16 activator protein (a chimera of the Tet repressor protein and VP16) is expressed and constitutively bound to the TetO binding sites but dissociates upon binding to tetracycline or its analog doxycycline. Therefore, the tRNA will be repressed by TetO/TetR-VP16 induced RNAP II transcription in medium lacking doxycycline and expressed when doxycycline is added. As shown in Figure S3 with β-galactosidase assays, transcription from the *CYC1-TetO* promoter decreases linearly with increasing concentrations of doxycycline in the range of 1 ng/mL to 100 ng/mL. The relative growth calculated from doubling times of strains containing wild type tRNA^Ser^-TetO, tRNA^Ser^_UGG_-TetO and tRNA^Ser^_UGG,G26A_-TetO when grown in media containing different concentrations of doxycyline is shown Figure 3B. In medium lacking doxycycline the wild type and tRNA^Ser^_UGG,G26A_ strains grow at a similar rate, while the tRNA^Ser^_UGG_ containing strain grows slightly slower. Increasing doxycycline does not decrease the growth of the wild type tRNA^Ser^-TetO strain. In contrast the growth of the tRNA^Ser^_UGG_-TetO strain decreased to ∼ 80% in medium containing 10 ng/mL doxycycline compared to growth in medium lacking doxycycline. Growth was virtually eliminated with 100 ng/mL doxycycline. A similar pattern of decreasing relative growth was seen with tRNA^Ser^_UGG_,_G26A_-TetO.

**Figure 3.**
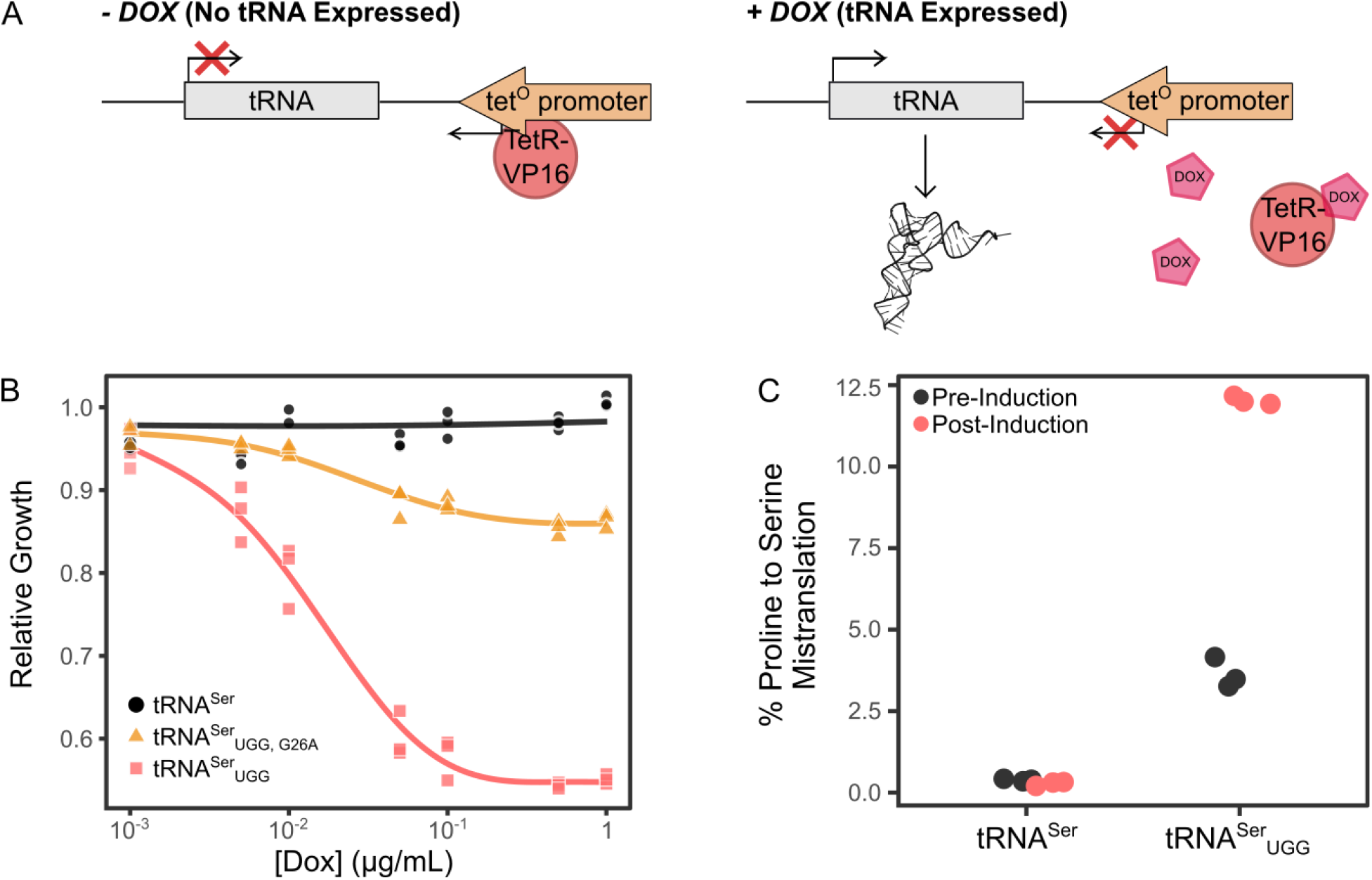
A doxycycline inducible tRNA expression system. **(A)** A model of the tRNA expression system with or without doxycycline. In the absence of doxycycline, TetR-VP16 mediated transcription from a downstream promoter containing 7 *tetO* binding sites interferes with RNAP III transcription of the tRNA. When doxycycline is added, TetR-VP16 dissociates from the promoter which allows RNAP III to transcribe the tRNA. **(B)** Yeast strain CY8652 constitutively expressing the TetR-VP16 protein and containing *LEU2* plasmids expressing either wild type tRNA^Ser^ or the mistranslating variants tRNA^Ser^_UGG,G26A_ or tRNA^Ser^_UGG_ were grown to saturation in medium lacking leucine without doxycycline. Strains were diluted to OD_600_ ∼ 0.1 in the same medium containing varying concentrations of doxycycline and grown for 24 hours at 30°C with agitation. OD_600_ was measured every 15 minutes and doubling time in minutes calculated from the growth curves. Doubling time was normalized to the doubling time of the wild type strain grown without doxycycline and used to calculate the relative growth of each strain at the different doxycycline concentrations. Each point represents a biological replicate (n = 3). **(C)** Mass spectrometry analysis of the cellular proteome was performed on yeast strain CY8652 constitutively expressing TetR-VP16 and containing a *LEU2* plasmid expressing either wild type tRNA^Ser^-TetO or the inducible tRNA^Ser^_UGG_-TetO. Cells were grown overnight in media lacking uracil and leucine, diluted to an OD_600_ of 0.05 in the same media and grown to an OD_600_ of ∼ 0.2 before they were treated with 10 μg/mL doxycycline. Aliquots were taken just before doxycycline addition (pre-Induction) and at 10 hours after (post-Induction) to determine proline to serine mistranslation frequency using mass spectrometry. Each point represents a biological replicate (n = 3).

We performed mass spectrometry to determine the maximum amount of mistranslation from tRNA^Ser^_UGG_-TetO before and after the addition of doxycycline. Prior to addition of doxycycline the mistranslation of serine at proline codons by tRNA^Ser^_UGG_ was 3.6% (Figure 3C). Ten hours after the addition of doxycycline, we observed 12% substitution of serine at proline codons.

To determine if the regulatory system was applicable to other tRNAs, we tested if a downstream TetR-VP16 activated promoter regulated mistranslation from an alanine tRNA variant. Since a G3:U70 base pair within the acceptor stem is the principal identity element for alanylation by AlaRS^36–39^, we replaced the cognate AGC anticodon of tRNA^Ala^ with UGG for proline (Figure 4A) to generate a mistranslating tRNA^Ala^. The tRNA^Ala^_UGG_ encoding gene was cloned precisely in place of tRNA^Ser^ to flank tRNA^Ala^_UGG_ gene with the tRNA^Ser^ 5’ and 3’ sequences (see Figure S4). Transformants were isolated and examined for growth in the presence or absence of doxycycline. tRNA^Ala^ with an alanine anticodon was cloned in the same setup and used as the control. Representative growth curves are shown in Figure 4B. When grown in the absence of doxycycline the strain containing tRNA^Ala^_UGG_ grew slower than the strain expressing tRNA^Ala^ (doubling times of 123 ± 3 and 92 ± 1 minutes from three biological replicates, respectively), indicative of mistranslation by tRNA^Ala^_UGG_ and again consistent with some leakiness of tRNA expression. In the presence of 1 μg/mL doxycycline, the doubling time of tRNA^Ala^_UGG_ increased to 194 ± 8 minutes, whereas tRNA^Ala^ was unchanged.

**Figure 4.**
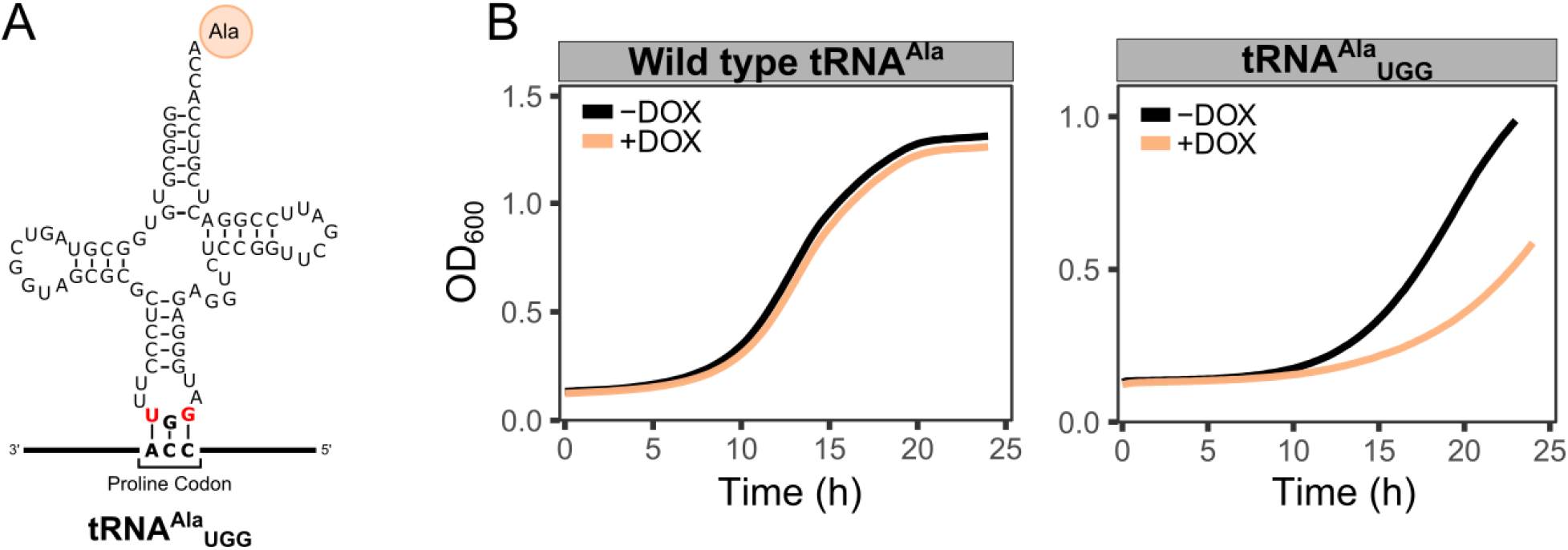
**(A)** Schematic of tRNA^Ala^_UGG_ which inserts alanine at proline codons. **(B)** Yeast strain CY8652 expressing either wild type tRNA^Ala^-TetO or tRNA^Ala^_UGG_-TetO were grown in medium lacking uracil and leucine, diluted to an OD_600_ of 0.1 in the same medium either with 1 μg/mL or without doxycycline and grown for 24 hours at 30°C with agitation. OD_600_ was measured every 15 minutes. One representative growth curve is shown for each condition.

### Impact of high levels of mistranslation on yeast cells

Next, we used the inducible system to investigate the effect of increasing tRNA derived mistranslation on growing cells. Yeast strains were inoculated into medium lacking doxycycline and grown for six hours before doxycycline was added to induce tRNA expression. As shown by the growth curves in Figure 5A, there was no difference in growth rate after doxycycline addition in the strain containing wild type tRNA^Ser^, whereas tRNA^Ser^_UGG_ began to slow growth four hours after doxycycline addition. The strain containing tRNA^Ser^_UGG_ stopped growing 15 hours after doxycycline addition, reaching an OD_600_ of approximately 2.0 compared to the uninduced tRNA^Ser^_UGG_ which reached an OD_600_ of greater than 5.0 at 15 hours.

**Figure 5.**
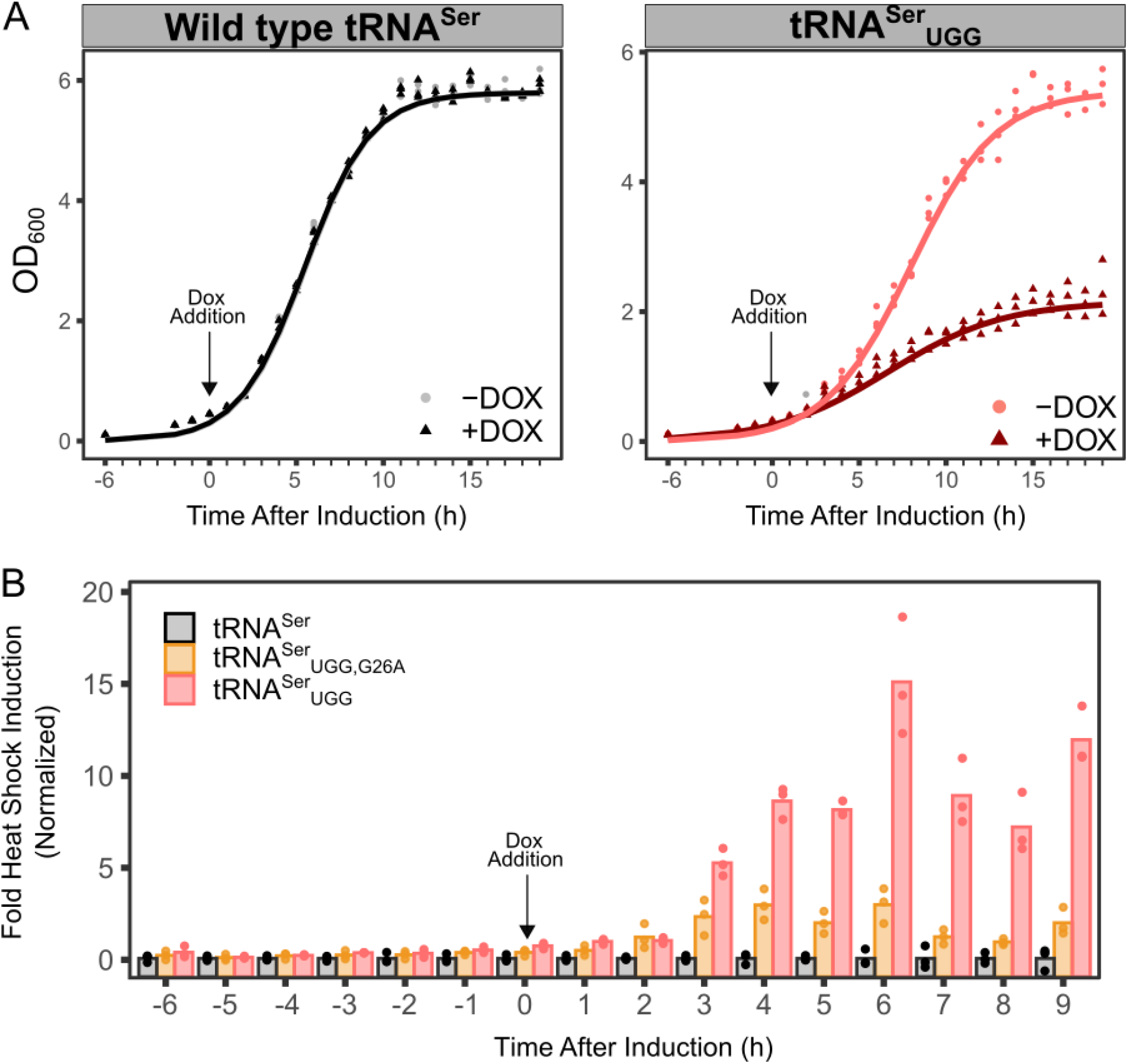
Effect of inducing high levels of tRNA derived mistranslation on cell growth and heat shock response. **(A)** Yeast strain CY8652 constitutively expressing the TetR-VP16 protein and containing *LEU2* plasmids expressing either wild type tRNA^Ser^-TetO or the mistranslating variant tRNA^Ser^_UGG_-TetO were grown to saturation in medium lacking leucine without doxycycline. Strains were diluted to OD_600_ ∼ 0.1 in the same medium and grown at 30°C with agitation. At OD_600_ of ∼ 0.3, cultures were either left to grow (-DOX) or treated with doxycycline at a concentration of 10 μg/mL (+DOX). OD_600_ was measured every hour. The arrow indicates the time of doxycycline addition. Each point represents a biological replicate (n = 3). **(B)** Heat shock response upon induction of the mistranslating tRNA variant. Yeast strain CY8652 constitutively expressing the TetR-VP16 protein and containing an *HSE-GFP* reporter plasmid with *LEU2* plasmids expressing either wild type tRNA^Ser^-TetO or the mistranslating variants tRNA^Ser^_UGG,G26A_-TetO or tRNA^Ser^_UGG_-TetO were grown to saturation in medium lacking leucine and histidine without doxycycline. Strains were diluted in the same medium to an OD_600_ of 0.1 and grown to an OD_600_ of ∼ 0.3 at 30°C before doxycycline was added to 10 μg/mL. Every hour, an aliquot of cells was removed from the culture, cell densities were normalized to OD_600_ of 0.1 and fluorescence was measured. Fold induction was calculated as the fluorescence signal relative to the strain expressing a wild type tRNA^Ser^ at each time point. The arrow indicates the time of doxycycline addition. Each point represents one biological replicate (n = 3).

Mis-made proteins arising from tRNA mediated mistranslation cause proteotoxic stress and induce a heat shock response in yeast^11,28,36,40^. Taking advantage of the inducible tRNA expression system, we investigated the dynamics of heat shock induction upon elevated expression of a mistranslating tRNA. Using a fluorescent GFP reporter driven by a promoter containing heat shock response elements, the extent of heat shock response was measured at various points after inducing either tRNA^Ser^_UGG_, which mistranslates at lethal levels, or tRNA^Ser^_UGG,G26A_, which mistranslates at levels that are tolerated by cells (Figure 5B). Three hours after doxycycline addition, GFP fluorescence increased 6.2-fold in the strain containing tRNA^Ser^_UGG_ and 3.3-fold for the less severe tRNA^Ser^_UGG,G26A_. The heat shock response reached a maximum increase of ∼16-fold and 4-fold at six hours after tRNA induction for tRNA^Ser^_UGG_ and tRNA^Ser^_UGG,G26A_, respectively.

To determine if cells resume growth after experiencing high levels of tRNA derived mistranslation, an aliquot of cells at various timepoints after doxycycline addition was washed in medium lacking doxycycline and spotted on solid medium lacking doxycycline (Figure 6A). Before adding doxycycline, strains containing tRNA^Ser^_UGG_ or tRNA^Ser^_UGG,G26A_ were viable. Viability decreased five hours after inducing tRNA expression with doxycycline for the strain expressing tRNA^Ser^_UGG_ and few viable cells remained after ten hours. In contrast, cells expressing the less severe tRNA^Ser^_UGG,G26A_ were still viable 20 hours after doxycycline addition. To determine if the high level of mistranslation causes cell death, cells were stained with propidium iodide at the same time points. Propidium iodide is a membrane-impermeable dye and only stains DNA in dead cells^41^. Interestingly, there was no increase in propidium iodide signal after increasing the expression of mistranslating tRNA^Ser^_UGG_ with the addition of doxycycline (Figure S5). This suggests that the mistranslation does not disrupt cellular membranes and the inability of the mistranslating cells to grow may not be due to typical cell death.

**Figure 6.**
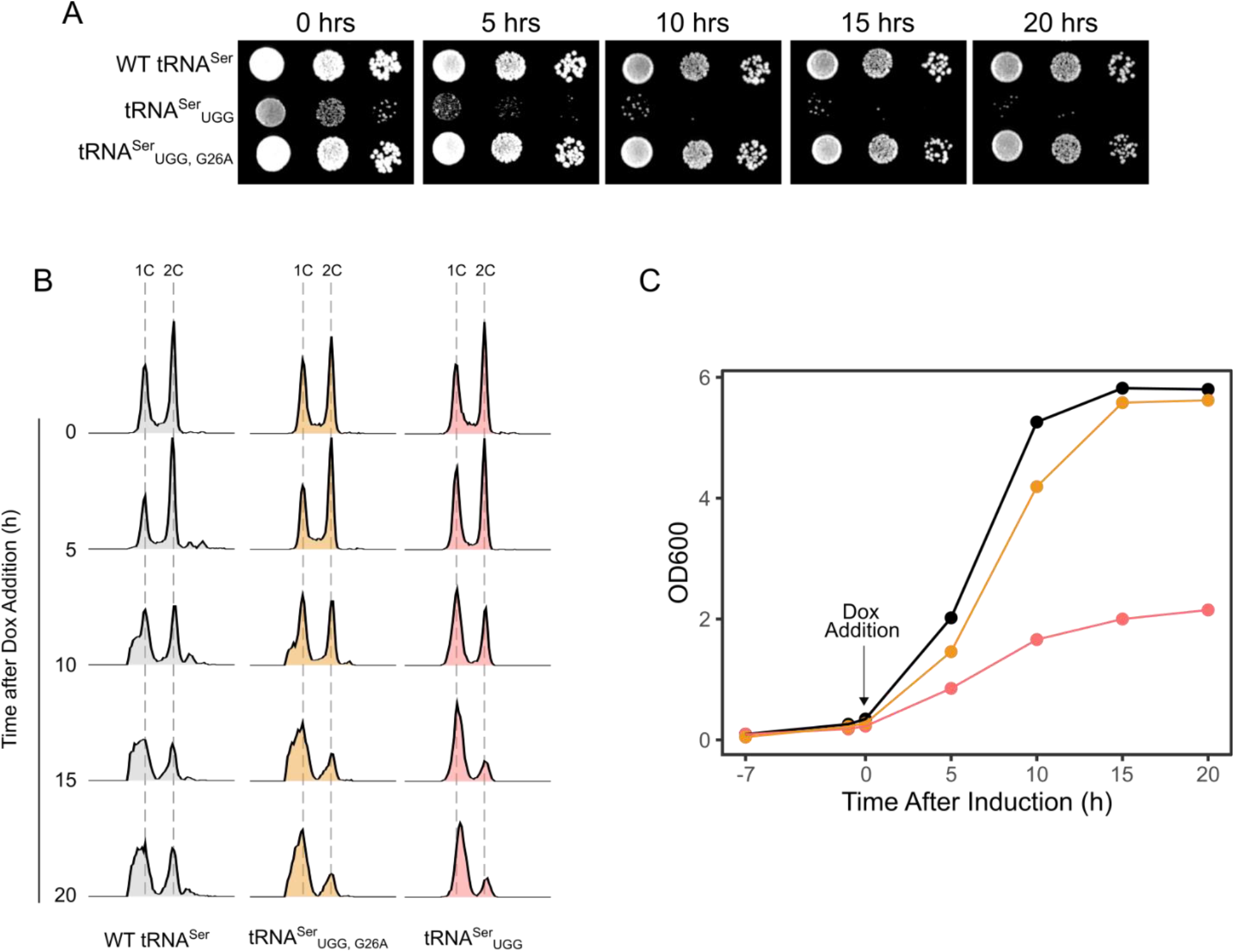
Growth is inhibited after high levels of tRNA based mistranslation. **(A)** Yeast strain CY8652 constitutively expressing the TetR-VP16 protein and containing *LEU2* plasmids expressing either wild type tRNA^Ser^-TetO or the mistranslating variant tRNA^Ser^_UGG_-TetO or tRNA^Ser^_UGG,G26A_-TetO were grown to saturation in medium lacking leucine without doxycycline. Strains were diluted to OD_600_ ∼ 0.1 in the same medium and grown at 30°C with agitation. At OD_600_ of ∼ 0.3, cultures were treated with doxycycline at a concentration of 10 μg/mL. At various time points after doxycycline addition, an equal number of cells of each strain were washed in medium lacking leucine without doxycycline and plated in 10-fold serial dilutions on the same medium lacking doxycycline and grown at 30°C. Plates were imaged after two days of growth. **(B)** Flow cytometric analysis of samples taken at various time points after doxycycline addition as in (A). Samples were taken at the times indicated in hours following doxycycline addition, prepared for analysis and stained with propidium iodide. Histograms represent ∼10,000 cells. Positions of cells with 1C and 2C DNA content are indicated on the x-axis, which reflects fluorescence intensity on a linear scale. The y-axis represents cell frequency and has been scaled to represent the percentage of the maximum bin contained in that graph. One representative biological replicate is shown. **(C)** Representative growth curve of strains used for flow cytometry analysis in (B). The arrow indicates the time of doxycycline addition.

We hypothesized that high levels of mistranslation might cause cells to arrest at a specific cell cycle stage. We performed flow cytometry after staining with propidium iodide at different time points after doxycycline addition to assess the proportion of cells in each stage of the cell cycle (Figure 6C). At 20 hours after doxycycline addition, strains expressing either tRNA^Ser^_UGG_ or tRNA^Ser^_UGG,G26A_ accumulated in G1 phase. In contrast, cells expressing wild type tRNA^Ser^ were approximately evenly split between G1 and G2 phases 20 hours after doxycycline addition. Both heat shock and the mis-incorporation of non-canonical amino acids also cause yeast to arrest at G1^42,43^. Differing from what we see with high levels of mistranslation, cells continue normally through the cell cycle after these stresses are removed^43^. We predict that the difference results from the inability of cells to sufficiently clear the extreme levels of mis-made protein caused by mistranslation to allow further growth.

From the cell cycle data, we noticed that in the more severe mistranslating strain expressing tRNA^Ser^_UGG_, the peaks corresponding to G1 and G2 DNA content shifted to the right, suggesting a possible change in cell size or shape. To determine if cell size and shape changes upon exposure to mistranslation, we imaged cells at various time points after doxycycline addition and measured the area of each cell as well as the ratio between the longest and shortest diameter (aspect ratio). There was no change in cell size or aspect ratio for strains expressing wild type tRNA^Ser^ across any of the time points. In contrast, cells containing the mistranslating tRNA^Ser^_UGG_ were larger than the control wild type tRNA^Ser^ cells (Figure 7A). Cells containing tRNA^Ser^_UGG_ and treated with doxycycline had greater mean area than untreated cells at both 8 and 24 hours after doxycycline addition. Cells containing tRNA^Ser^_UGG_ also shifted towards a greater aspect ratio than control cells containing wild type tRNA^Ser^ (Figure 7B). This difference was exaggerated 24 hours after doxycycline addition. As shown by the representative cell images in Figure 7C, and in agreement with the increased aspect ratio, mistranslation results in cells with a more oblong shape.

**Figure 7.**
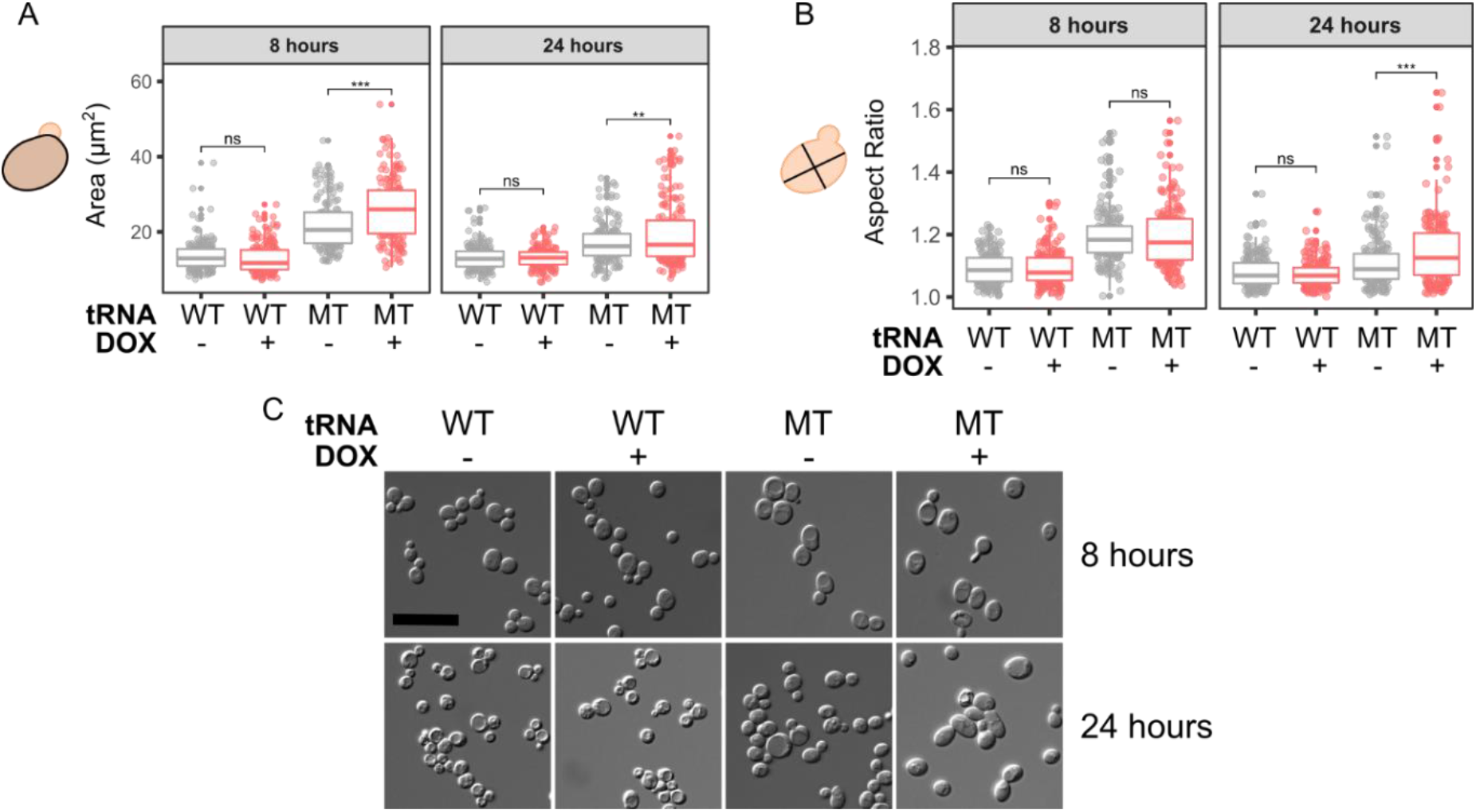
Cell size and shape changes upon exposure to mistranslation. Images of cells either containing an inducible wild type tRNA^Ser^ or mistranslating tRNA^Ser^_UGG_ were taken at various time points after doxycycline addition. Strains expressing either wild type tRNA^Ser^-TetO or tRNA^Ser^_UGG_-TetO were diluted to an OD_600_ of 0.1 in media lacking doxycycline. At an OD_600_ of ∼ 0.3, doxycycline was added to a final concentration of 10 μg/mL. Samples were taken at eight and 24 hours after doxycycline addition and imaged at 63x magnification using DIC. **A**. Area and **B**. aspect ratio (ratio between the longest and short diameter of each cell) were measured for 80 cells at each time point. Each point represents one cell and stars indicate significant differences (Welch’s T-test; ** *p* < 0.001, *** *p* < 0.0001; ns – not statistically different). **C**. Representative images of cells from each strain with and without doxycycline treatment at each time point. Scale bar represents 20 μm.

## CONCLUSIONS

We have created regulated tRNA expression systems where tRNA expression is repressed by a downstream RNAP II promoter directed towards the tRNA encoding gene. Since repression required a transcriptional activator and occurred only when the downstream promoter was actively transcribed, we conclude that transcription from the RNAP II promoter is required rather than just activator binding. We predict that repression is due to a readthrough transcript based on the proximity of the promoter to the tRNA encoding gene. If the active promoter were simply leading to more open chromatin, one would expect to see increased RNAP III expression rather than repression.

The tetO system allowed for titratable expression of an otherwise toxic mistranslating tRNA. When not repressed, tRNA^Ser^_UGG_ incorporated serine at proline codons at a maximum frequency of 12%. In preliminary studies we have observed that the toxicity resulting from misincorporation due to two different mistranslating tRNA variants is additive. By placing two different mistranslating tRNAs under regulation it should be possible to mistranslate codons for different amino acids at a high level simultaneously.

Our system has applications in regulating the expression of a variety of tRNAs in addition to mistranslating tRNAs. In genetic code expansion applications, the orthogonal tRNA synthetase is normally placed under regulated control to mediate the toxic effects of mis-incorporation at stop codons. Using our system, the tRNA can also be regulated, allowing prior cell growth and enhanced expression of modified protein. Ambiguous decoding has applications in biosafety because it allows expression of dangerous products without the threat of horizontal gene transfer^44^. Allowing high level expression of the required tRNA will further the utility of this approach by increasing expression of the product. Endogenous wild type tRNAs can also be regulated by this system to investigate the effects of modulating cellular tRNA concentrations on translation and the proteome. Variations of this inducible tRNA system are applicable to other eukaryotes including multicellular model organisms such as *Drosophila melanogaster* and *Caenorhabditis elegans*.

Using the inducible tRNA system, we investigated the effects of high levels of mistranslation on cell growth. A heat shock response was detectable ∼ 3 hours after doxycycline addition to induce greater expression of the mistranslating tRNA. Growth rate decreased one hour later. Cells exposed to high levels of mistranslation arrest in G1 in an irreversible manner and display an altered size and shape.

tRNA expression was not fully repressed in either the Gal4 regulated or the tetracycline regulated system. Moving the promoter closer to the tRNA or adding an additional flanking promoter did not appreciably alter repression. It may be possible to increase repression by increasing RNAP II promoter strength with additional activator binding sites, a stronger activator and/or increased activator expression. Flanking sequence could also be analyzed for sequences that impede passage of RNAP II into the tRNA encoding gene. In this regard we found that the native sequences flanking tRNA^Ala^ did not permit repression. It might also be possible to obtain a wider range of tRNA expression levels using a RNAP II promoter with more gradual changes in its expression, such as the β-estradiol promoter^45,46^. For tRNAs where leaky expression still results in lethality, expression can be facilitated by incorporating a secondary mutation such as G26A that partially destabilizes the tRNA.

## MATERIALS AND METHODS

### Yeast strains and growth

Wild type haploid yeast strains are derivatives of BY4742 (***MAT****α his3Δ1 leu2Δ0 lys2Δ0 ura3Δ0*)^47^. The haploid strain CY8652 (***MAT****α his3Δ1 leu2Δ0 lys2Δ0 ura3Δ0 tTA*-URA3)* containing the tet “off” activator (tTA*) marked with *URA3* was derived from R1158^35^ after crossing with BY4741 and sporulation. The yeast two hybrid strain PJ69-4a (***MAT****a trp1-901 leu2-3,112 ura3-52 his3-200 gal4Δ gal80Δ LYS2::GAL1-HIS3 GAL2::ADE2 met2::GAL7-lacZ*) was described in James *et al*. (1996)^48^.

Yeast strains were grown at 30°C in yeast peptone medium or in synthetic medium supplemented with nitrogenous bases and amino acids containing 2% glucose, 2% galactose or 2% raffinose as indicated. Transformations were performed using the lithium acetate method as performed in Berg *et al*. (2018)^49^. Growth curves were generated by diluting saturated cultures to OD_600_ ∼ 0.1 in synthetic medium and incubating at 30°C. OD_600_ was measured every 15 minutes for 24 hours in a BioTek Epoch 2 microplate spectrophotometer or by hand every hour. Doubling time was calculated using the R package “growthcurver”^50^.

### DNA constructs

*SUP17* (tRNA^Ser^; pCB3076), *sup17(UGG)* (tRNA^Ser^_UGG_; pCB3082) and *sup17(UGG)-G26A* (tRNA^Ser^_UGG, G26A_; pCB4023), including approximately 300 base pairs of 5’ and 3’ sequence, in YCplac33 have been described previously^15,28^. The *GAL1* promoter was amplified from yeast genomic DNA using primers 4588-1/XH5103 and ligated into pGEM-Teasy (Promega) to create pCB4554. *GAL1* promoter was cut with *Hin*dIII and ligated into pCB4023 to put the *GAL1* promoter upstream of *sup17(UGG)-G26A* to create pCB4568.*GAL1* promoter was cut with *Eco*RI and ligated into pCB4023 to put the *GAL1* promoter downstream of *sup17(UGG)-G26A* to create pCB4566. To create the construct with two *GAL1* promoters, the *GAL1* promoter was cut with *Hin*dIII and ligated into pCB4566 to create pCB4851.

The synthetic *HIS3* promoter with five Gal4 binding sites (pCB859; *his3*-G4) was previously described in Brandl *et al*. (1993)^33^. The promoter was amplified with primers XI6247/XI6248 and cloned into pGEM-Teasy (Promega) to create pCB4594. The tRNAs *SUP17* (pCB3076), *sup17(UGG)* (pCB3082) or *sup17(UGG)-G26A* (pCB4023) were amplified with UG5953/UG5954 and cloned into pGEM-Teasy (Promega) to create pCB4652, pCB4603 and pCB4593, respectively. The synthetic *GAL* promoter was cut with *Kpn*I-*Not*I and cloned downstream of each tRNA cut *Not*I-*Hin*dIII in vector YCplac33 cut *Hin*dIII-*Kpn*I to create pCB4657, pCB4612 and pCB4598, respectively.

Constructs with different spacing between the tRNA 3’ end and promoter were created by amplifying *sup17(UGG)* (pCB3082) with upstream primer UG5953 and downstream primers YA9566 (100 bp) or YA9567 (200 bp) and ligated into pGEM-Teasy (Promega) to create pCB4633 and pCB4634, respectively. Each tRNA construct was cut with *Not*I-*Hin*dIII and cloned into a vector containing the synthetic *GAL* promoter to create pCB4660 and pCB4661, respectively.

The minimal *CYC1* promoter with seven tetracycline binding sites was amplified from the yeast Tet-Promoters collection^35^ with YG4866/YG4867 and cloned into pGEM-Teasy (Promega) to create pCB4695. The TetO promoter was cut *Bam*HI-*Not*I and cloned with *SUP17, sup17(UGG)* or *sup17(UGG)-G26A* cut *Hin*dIII-*Not*I into YCplac111 cut *Bam*HI-*Hin*dIII to create pCB4699, pCB4700 and pCB4701, respectively.

The Tet promoter was amplified from the yeast Tet-Promoters collection^35^ with primers YG4866/YG4868 and cloned as a *Bam*HI-*Hin*dIII fragment into the *LEU2* centromeric plasmid YCplac87 to give the a *his3-lacZ* fusion reporter pCB4705^33^.

The inducible wild type tRNA^Ala^ and mistranslating tRNA^Ala^_UGG_ were synthesized as a GeneString (ThermoFisher; Figure S4) and cloned as *Hin*dIII-*Not*I fragments into pCB4699 adjacent to the Tet promoter to create pCB4787 and pCB4776, respectively.

The centromeric plasmid containing the *HSE-eGFP* was kindly provided by Onn Brandman (Stanford University) ^51^. The plasmid expressing the *GAL4* DNA binding domain was kindly provided by Ivan Sadowski^52^.

### Mass spectrometry

Liquid chromatography tandem mass spectrometry was performed on strains expressing mistranslating tRNA variants to identify mistranslation. For strains containing the *GAL1* regulated tRNA constructs, starter cultures of each strain were grown to saturation in medium lacking uracil and containing 2% galactose, diluted 1:20 in the same media and grown for 18 h at 30°C. For the strains containing the *tetO* regulated tRNA constructs, starter cultures of each strain were grown to saturation in media lacking uracil and leucine, diluted to an OD_600_ of 0.05 in the same media and grown to an OD_600_ of ∼ 2 before 10 μg/mL doxycycline was added. Preparation of cell lysates, protein reduction and alkylation were performed as described in Berg *et al*. (2019b)^15^.

Robotic purification and digestion of proteins into peptides were performed on the KingFisher™ Flex using LysC and the R2-P1 method as described in Leutert *et al*. (2019)^53^. Peptides were analyzed on a hybrid quadrupole orbitrap mass spectrometer (Orbitrap Exploris 480; Thermo Fisher Scientific) equipped with an Easy1200 nanoLC system (Thermo Fisher Scientific). Peptide samples were resuspended in 4% acetonitrile, 3% formic acid and loaded onto a 100 μm ID × 3 cm precolumn packed with Reprosil C18 3 μm beads (Dr. Maisch GmbH) and separated by reverse-phase chromatography on a 100 μm ID × 30 cm analytical column packed with Reprosil C18 1.9 μm beads (Dr. Maisch GmbH) housed into a column heater set at 50°C.

Peptides were separated using a gradient of 5-30% acetonitrile in 0.125% formic acid at 400 nL/min over 95 min and online analyzed by tandem mass spectrometry with a total 120 minute acquisition time. The mass spectrometer was operated in data-dependent acquisition mode with a defined cycle time of 3 seconds. For each cycle one full mass spectrometry (MS) scan was acquired from 350 to 1200 m/z at 120,000 resolution with a fill target of 3E6 ions and automated calculation of injection time. The most abundant ions from the full MS scan were selected for fragmentation using 2 m/z precursor isolation window and beam-type collisional-activation dissociation (HCD) with 30% normalized collision energy. MS/MS spectra were acquired at 15,000 resolution by setting the AGC target to standard and injection time to automated mode. Fragmented precursors were dynamically excluded from selection for 60 seconds.

MS/MS spectra were searched against the *S. cerevisiae* protein sequence database (downloaded from the Saccharomyces Genome Database resource in 2014) using Comet (release 2015.01)^54^. The precursor mass tolerance was set to 50 ppm. Constant modification of cysteine carbamidomethylation (57.0215 Da) and variable modification of methionine oxidation (15.9949 Da) and proline to serine (−10.0207 Da) were used for all searches. A maximum of two of each variable modification were allowed per peptide. Search results were filtered to a 1% false discovery rate at the peptide spectrum match level using Percolator^55^. The mistranslation frequency was calculated using the unique mistranslated peptides for which the non-mistranslated sibling peptide was also observed. The frequency is defined as the counts of mistranslated peptides, where serine was inserted for proline divided by the counts of all peptides containing proline or the mistranslated serine and expressed as a percentage. The mass spectrometry proteomics data have been deposited to the ProteomeXchange Consortium via the PRIDE^56^ partner repository with the dataset identifier PXD028496.

### B-Galactosidase assay

Yeast strain CY8652 containing pCB4705 was grown to stationary phase, diluted 10-fold into media containing various concentrations of doxycycline and grown for eight hours at 30°C. β-galactosidase units were determined using o-nitrophenyl-β-galactoside as substrate with values normalized to cell densities as described by Ausubel *et al*. (1989)^57^.

### Cell viability assay

Yeast strain CY8652 containing an inducible tRNA was grown to saturation in medium lacking leucine and uracil, diluted to an OD_600_ of 0.1 in the same media and grown for six hours before doxycycline was added to a final concentration of 10 μg/mL. Cell viability was assessed using propidium iodide as described in Chadwick *et al*. (2016)^41^. Briefly, cells were washed in phosphate-buffered saline (PBS) then resuspended in PBS containing 5 μg/mL propidium iodide (Invitrogen). A sample was boiled before resuspension in propidium iodide as a positive control and an unstained sample was used as the negative control. Samples were incubated at room temperature for 10 minutes before imaging on a Gel Doc system (Bio-Rad). The OD_600_ of each sample was determined using a BioTek Epoch 2 microplate reader.

### Flow cytometry

Yeast strain CY8652 containing an inducible tRNA were grown to saturation in medium lacking leucine and uracil, diluted to an OD_600_ of 0.1 in the same media and grown for six hours before doxycycline was added to a final concentration of 10 μg/mL. An aliquot of ∼ 10^7^ cells were harvested every five hours for 20 hours after doxycycline addition. Cells were prepared for flow cytometry as described in Bellay *et al*. (2011)^58^. Briefly, cells were fixed in 70% ethanol for at least 15 minutes at room temperature, washed with water and incubated sequentially in 0.2 mg/mL RNase A (Sigma) for 2 hours at 37°C and then 2 mg/mL proteinase K (BioBasic) for 40 minutes at 50°C. Cells were resuspended in FACS buffer (200 mM Tris, pH 7.5, 200 mM NaCl and 78 mM MgCl_2_) and stained with 5 μg/mL propidium iodide (Invitrogen) in FACS buffer. The samples were briefly sonicated and analyzed using a BD FACSCelesta flow cytometer (Becton Dickinson Biosciences). Data was analyzed using FlowJo Flow Cytometry Analysis software and plotted on a linear scale.

### Fluorescence heat shock assay

Yeast strain CY8652 expressing TetR-VP16 and containing the *HSE-GFP* reporter and inducible tRNA were grown to saturation in medium lacking leucine, uracil and histidine, diluted to an OD_600_ of 0.1 in the same medium. Every hour cell densities were normalized to OD_600_ of 0.1 and fluorescence measured with a BioTek Synergy H1 microplate reader at an emission wavelength of 528 nm. Doxycycline was added six hours after initial dilution to a final concentration of 10 μg/mL. At each time point, relative fluorescence units were calculated by subtracting background signal from a yeast strain lacking the HSE-GFP reporter. Fold induction was calculated by normalizing to the strain expressing wild type tRNA_Ser_.

### Microscopy

Yeast strain CY8652 containing either wild type tRNA^Ser^-TetO or the mistranslating tRNA^Ser^_UGG_-TetO were diluted to an OD_600_ of 0.1 in medium lacking uracil and leucine and grown for 6 hours at 30°C. After six hours, cells were either treated with 10 μg/mL doxycycline or left to grow. Cells were imaged at 8 and 24 hours after addition of doxycycline at 63x magnification using DIC on a Zeiss Axio Imager Z1 Fluorescent microscope using ZEN Blue Pro software (Zeiss Inc.). Cell size and aspect ratio was quantified by outlining living, non-budding cells in ImageJ^59^.

## ACKNOWLEDGMENTS

We thank Ricard Rodriguez-Mias for assisting with the mass spectrometry and maintaining the instruments, Ivan Sadowski for providing Gal4 expressing plasmids, Philip James for the PJ69-4a strain, Tina Sing for assistance with analysis of FACS data and the Biotron Integrated Microscopy facility staff for their help with the yeast microscopy.

## FUNDING

This work was supported by the Natural Sciences and Engineering Research Council of Canada (NSERC) [RGPIN-2015-04394 to C.J.B.] and generous donations from Graham Wright and James Robertson to M.D.B. PL is supported by an NSERC Discovery Grant [RGPIN-2015-06300] and a Canadian Foundation for Innovation (CFI) John R. Evans Leader Fund Grant [65183]. Mass spectrometry work was supported by a research grant from the Keck Foundation, NIH grant R35 GM119536 and associated instrumentation supplement (to J.V.). M.D.B. holds an NSERC Alexander Graham Bell Canada Graduate Scholarship (CGS-D). J.I. holds an NSERC Alexander Graham Bell Canada Graduate Scholarship (PGS-D).

## Notes

### Competing Interest Statement

The authors have declared no competing interest.

