## Supplemental Material for "Regulating expression of mistranslating tRNAs by readthrough RNA polymerase II transcription"

### Supplemental Tables

**Table S1.** Oligonucleotides used in this study.

| Name | Sequence | Description |
|---|---|---|
| 4588-1 | CCCGGATCCGACGTTAAAGTATAGAGGT | *GAL1pr* |
| XH5103 | CCCAAGCTTTAATACGCTTAACTGCTCATTG | *GAL1pr* |
| XI6247 | TTTGGTACCCCTTTAGCTTCTCGACGTGG | *HIS3pr* |
| XI6248 | TTTGCGGCCGCTCTTTGCCTTCGTTTATCTTGCC | *HIS3pr* |
| UG5953 | TCTAAGCTTCGGACGATTGCCAACCGCCGAA | *SUP17* |
| UG5954 | CTGCAGAATTCCGCGGAAATTAGCACGGCC | *SUP17* |
| YA9566 | AAAGCGGCCGCGAACCGTGGCTAATAGG | *SUP17* |
| YA9567 | TTTGCGGCCGCTTTCCCCGAGAAAGCAACC | *SUP17* |
| YG4866 | TTTGGATCCGGGATCAACTCGAGTAATTCG | *TetOpr* |
| YG4867 | GGGGCGGCCGCGTAATTTAGTGTGTGTATTTGTG | *TetOpr* |
| YG4868 | CCCAAGCTTTCATTAGTAATTTAGTGTGTGTATTTGTG | *TetOpr* |

### Supplemental Figures

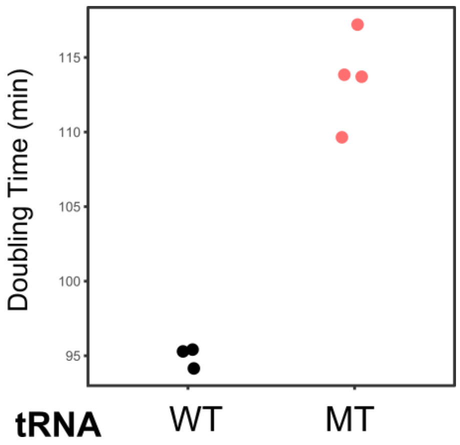


**Figure S1.** Expressing the *GAL4* DNA binding domain alone does not inhibit the toxicity due to $tRNA^{Ser}_{UGG,\ G26A}$ with downstream *HIS3*$^{5XGAL4}$ promoter. Yeast strain PJ69-4a (*gal4Δ gal80Δ*) expressing the *GAL4* DNA binding domain (pKW21) and either wild type $tRNA^{Ser}$ regulated by a downstream *HIS3*$^{5XGAL4}$ promoter (WT) or mistranslating $tRNA^{Ser}_{UGG,\ G26A}$ (MT) regulated by a downstream *HIS3*$^{5XGAL4}$ promoter, were grown to stationary phase in minimal medium. Strains were diluted to an $OD_{600}$ of 0.1 and grown for 24 hours at 30°C with agitation. $OD_{600}$ was measured every 15 minutes and doubling time in minutes was calculated from the growth curves.

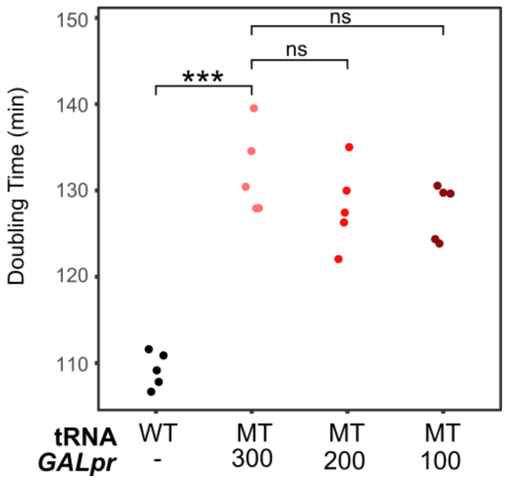


**Figure S2.** Distance between tRNA 3' end and *GAL1* promoter does not affect toxicity in galactose. Strains expressing wild-type $tRNA^{Ser}$ or mistranslating $tRNA^{Ser}_{UGG}$ variants with the synthetic *GAL1* promoter placed 300 bp, 200 bp or 100 bp from the 3' end of the tRNA were grown to stationary phase overnight in media lacking uracil and containing galactose. Strains were diluted to an $OD_{600}$ of 0.1 and grown for 24 hours at 30°C with agitation in media lacking uracil and containing either galactose or glucose as indicated. $OD_{600}$ was measured every 15 minutes and doubling time in minutes was calculated from the growth curves. Stars indicate significant differences (Welch's T-test, *** $P < 0.0005$) and ns indicates the differences was not statistically significant.

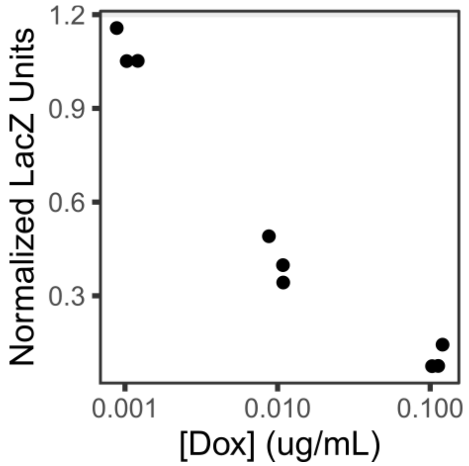


**Figure S3.** Effect of doxycycline concentration on transcription from *tetO* promoter. CY8652 containing the TetR-VP16 activator and *tetO*-LacZ reporter were grown to stationary phase in medium lacking uracil and leucine. Strains were diluted in the same medium containing various concentrations of doxycycline and grown for eight hours. B-galactosidase activity was determined and normalized to the strains grown in the absence of doxycycline. Each point represents one biological replicate.

AAGCTTC GGACGATTGC CAACCGCCGA AAAGGTTCAA GCCAAGAACA AAAAGTAGAG
AGAATACCCC AACAATAGCA TTTGATGGCT AAACGGCAGT GCCCGTTAAT ACTATTCAAA
TTATACCCCG CATTAATAAT TCCTGTCCCT TTAGTCTCGT CCTTCAGCCG TATAGCCGCA
AAACTTCGTT CAATGATTTC ATGCCATCAT TACACCGCGA GTCGTGGGTG CATTGTAGTT
TATTACGAAA AGTATCTACA ATACTTGCTT AAATAACCTA CATTGTTTTA **GGGCGTGTGG**
**CGTAGTCGGT AGCGCGCTCC CTTAGCATGG GAGAGGTCTC CGGTTCGATT CCGGACTCGT**
**CCA**TTATTTT TTTATTTTTA TTTTTTTTTT ATCGCTTACT GATTATCAGA TATCTTCGAC
AGCCTCACAC AGATTGGTGA TCACGCACCC ATAATCATTT CTTCGGGCAT GCTCCTATTA
GCCACGGTTC GCAGAATAAT CTGCGCGATG TTATTCACCA AGACCGTGGA GTCTCCTTTT
CGGTGGGATC CGCTTATATC CGTATGCGTC GGTTGCTTTC TCGGGGAAAG GAAAAGGAGA
AAGCCCGGAT AACACGACAC AAGAGTCATT CGTTATGCAC GGACACAGGG AATCGTGCGG
AGCCGGGGAT AAGAGGCCGT GCTAATTTCC GCGGAAGCGG CCGC

**Figure S4.** Inducible alanine mistranslating tRNA with $tRNA^{Ser}$ flanking sequence. Bolded sequence represents the alanine tRNA. Underlined sequence is the restriction sites used to clone the construct into the yeast expression plasmid.

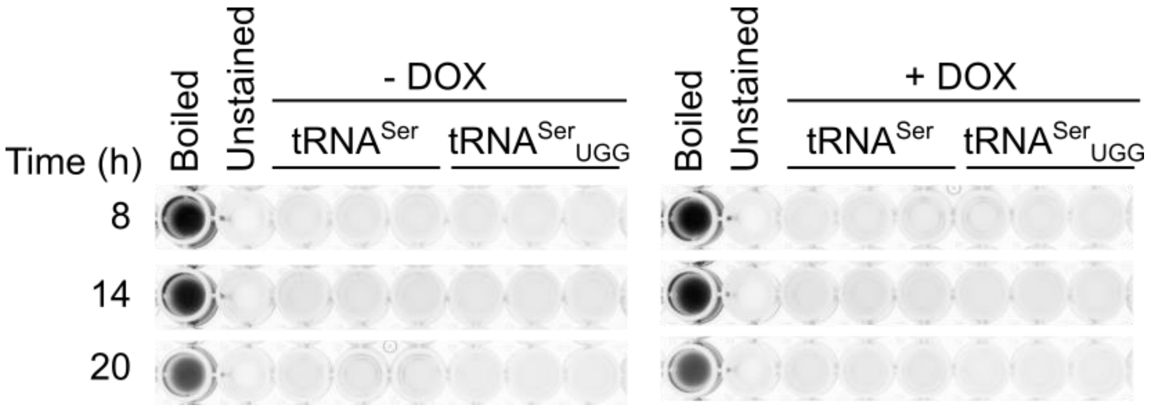


**Figure S5.** Inducing high levels of mistranslation does not increase propidium iodide staining. Strain CY8652 containing either $tRNA^{Ser}$ or the mistranslating $tRNA^{Ser}_{UGG}$ regulated by the *tetO* promoter were grown to stationary phase in medium lacking uracil and leucine. Strains were diluted in the same medium to an $OD_{600}$ of 0.1 and grown for 6 hours before doxycycline was added to a final concentration of 10 µg/mL. At various time points after doxycycline addition, aliquots of cells were stained with propidium iodide and imaged using a UV transilluminator. Percentage of cells alive was calculated relative to the boiled sample.
